# Deficiency in transmitter release triggers homeostatic transcriptional changes that increase presynaptic excitability

**DOI:** 10.1101/2024.08.30.610173

**Authors:** Caroline A. Cypranowska, Dariya Bakshinska, Maya Feldthouse, Yoon Gi Choi, Rachel Li, Zachary L. Newman, Ehud Y. Isacoff

## Abstract

Weakening of synaptic transmission at the *Drosophila* larval neuromuscular junction triggers two forms of homeostatic compensation, one that increases the probability of glutamate release per action potential (*P_r_*) and another that increases motoneuron (MN) activity. We investigated the molecular changes in MNs that underlie the increase in MN activity. RNA-seq analysis on MNs whose glutamate release is weakened by knockdown of components of the MN transmitter release machinery reveals a reduction in expression of a group of genes that encode potassium channels and their positive modulators. These results identify a mechanism of compensation for weakened synaptic transmission by MNs, which engages a transcriptional program in those cells to increase firing and, thereby, ensure sufficient locomotory drive.

## Introduction

To ensure reliable circuit function, the nervous system possesses homeostatic mechanisms that adjust neurotransmitter release to match postsynaptic sensitivity to transmitter, maintain a balance between excitatory and inhibitory inputs, restore total synaptic excitation to a transmission point following plasticity changes at a subset of synapses, while maintaining the newly differentiated synaptic weights, and regulate action potential (AP) firing to return to a firing set point following perturbation in order to ensure proper circuit output^1–3^. In the *Drosophila* larval neuromuscular junction (NMJ), pharmacological or genetic reduction of the excitatory postsynaptic current (EPSC) through the GluRII glutamate-gated channel is compensated for by an increase in the amount of per AP glutamate release by type I MNs^4–7^. However, this presynaptic homeostatic plasticity (PHP) occurs only at one of the two type I MNs, which converge onto each muscle: the type Ib MN, not at the Is MN, and the Ib compensation is incomplete^5,6^. Given that the type Ib MN generates a smaller excitatory postsynaptic potential (EPSP) than does the type Ib MN^8,9^ and given the incomplete compensation of the Ib input, PHP only provides incomplete compensation for postsynaptic weakening in the context of the entire MN-muscle unit.

The *Drosophila* larval locomotory circuit has a second mechanism for compensation that appears to make up the difference. Mutation of the voltage-gated K^+^ channel EAG that blocks calmodulin-binding reduces AP-evoked glutamate release and increases type I MN excitability^10^, suggesting a mechanism of firing compensation for synaptic weakening. This observation is complicated by the fact that the mutation is in a K^+^ channel that itself controls excitability. However, in support of this model, we recently observed that either postsynaptic weakening (due to mutation in the postsynaptic GluRII receptor) or presynaptic weakening (due to knockdown of Cac, the voltage-gated Ca^2+^ channel that couples the AP to release, or of vesicle priming proteins Rbp or Unc13) increases type I MN activity and preserves near normal locomotion^11^. The molecular mechanism of this MN firing compensation is unknown.

To probe the molecular mechanism of MN firing compensation, we examined the effect on gene expression of knockdown of the same presynaptic release components. We first confirmed that type I MN selective RNAi knockdown of Rbp or Unc-13 severely reduces AP-evoked glutamate release. RNA-seq analysis revealed that these knockdowns induce changes in expression of genes encoding release ion channels, machinery proteins, and neuropeptides. Among these, a group of seven genes, which encode either voltage-gated K^+^ (K_v_) channels or their positive modulators and auxiliary subunits stood out. Expression of all of these was reduced in the *Rbp* and *unc-13* knockdowns, consistent with increased excitability. We propose that weakening of transmitter release from MNs triggers the coordinate reduction in expression of a homeostatic transcriptional module that normally suppresses AP firing to ensure constancy in circuit output and behavior.

## Results

Chronic reduction of glutamate release from type I MNs in the *Drosophila* larval NMJ elicits a compensatory increase in presynaptic firing to maintain circuit output^10^. To understand the cell autonomous molecular mechanisms in MNs that mediate this compensation, performed RNA-seq on type I MNs whose glutamate release is compromised by knockdown of components of the presynaptic transmitter release machinery. To selectively perturb transmitter release in these neurons, we utilized OK6, a type I MN-specific Gal4 driver^13^, to drive expression of RNAi constructs targeting: (i) RIM-binding protein (Rbp), which connects release sites to the scaffolding protein Bruchpilot (Brp), (ii) Unc-13, a regulator of synaptic vesicle priming and (iii) Cacophany (Cac), the voltage-gated Ca^2+^ channel that couples the AP to transmitter release^14,15–18^. Synaptic transmission was imaged in the dissected preparation, where nerve bundles containing Ib and Is MN axons are stimulated electrically. We used spinning disk confocal microscopy to visualize the Ca^2+^ component of cation influx through the GluRII receptors that generates the excitatory postsynaptic current. To do this, we utilized SynapGCaMP6f, which targets the postsynaptic density and enables optical quantal analysis^5,19,20^. We applied our recently developed QuaSOR, <u>Qua</u>ntal <u>S</u>ynaptic <u>O</u>ptical <u>R</u>econstruction, to enhance the spatial resolution of quantal transmission imaging^21^. In brief, QuaSOR applies 2D Gaussian fits to transmission events to more precisely localize their origin, enabling discrimination of concurrent release events even in areas of densely packed synapses. As an optical measure of quantal content, we measured quantal density, the number of evoked transmission events per stimulus per unit area of postsynaptic membrane^5^. In type Ib synapses, MN knockdowns of *Rbp*, *unc-13* or *cac* reduced quantal density by ∼70-85% (**Figure 1**). To focus on perturbations that selectively weaken transmission, we focused on knockdown of *Rbp* and *unc-13* and set aside *cac*, whose effect on calcium influx may alter additional processes in the cell beyond transmitter release.

**Figure 1.**
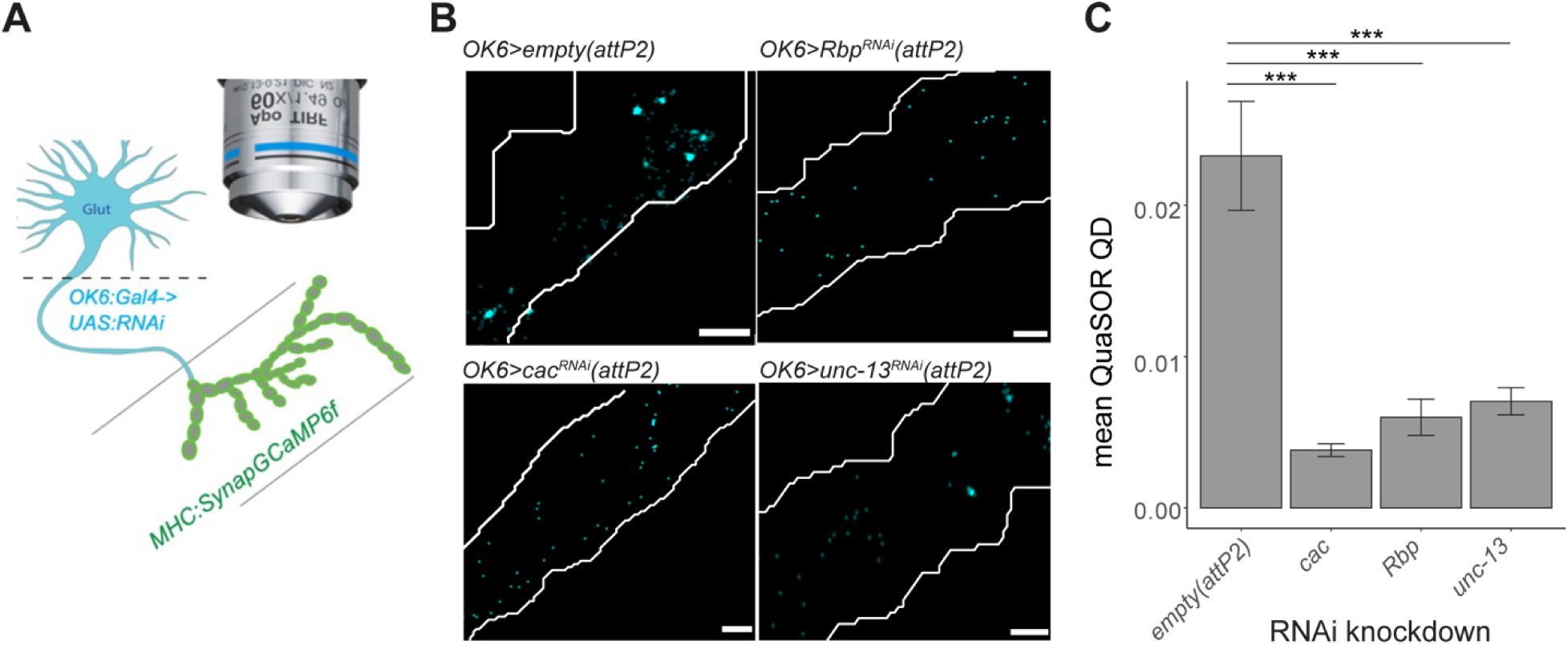
Weakened transmitter release from type Ib motor neurons by RNAi knockdown of components of the presynaptic transmitter release machinery. **A)** Quantal of imaging synaptic transmission from Ib MNs by SynapGCaMP6f expressed in larval body wall muscle under MHC promoter (green) and OK6:Gal4 driving expression of UAS-RNAi in MNs (blue). **B, C)** Cumulative transmission maps (**B**) and quantal density (average number of events per unit area of muscle) determined following QuaSOR analysis (**C**) in control (*attP2 (empty)*, n_Ib_ = 7) and UAS-RNAi for Rbp (n_Ib_ = 6), Unc13 (n_Ib_ = 5) or Cac (n_Ib_ = 5), each inserted at the *attP2* site. Scale bar: 2 μm. C) Quantal density values are then mean ± S.E.M. (One-tail Student’s t-test; *** p < 001).

### Knockdown of RSSP components in type I MNs induces broad changes in gene expression

To understand the molecular alterations that occur in type I MNs when synaptic transmission to muscle is profoundly compromised, we devised a scheme to sequence the transcriptomes of type I MNs, which carry the main excitatory drive to locomotory muscles in *Drosophila*, and to compare controls to animals in which one of the components of the presynaptic release machinery is knocked down exclusively in the type I MNs. We isolated type I MNs by expressing a nuclear mCherry reporter under the type I MN-specific *OK6-Gal4* line, dissecting the ventral nerve cords, dissociating the cells and capturing mCherry-expressing cells by fluorescence-activated cell sorting (FACS) (**Figure 2A**). We extracted the RNA from this purified population of cells and prepared sequencing libraries from pooled sorted cell samples from control animals (*attP2*) and from animals that expressed the RNAi construct for either unc-13 (*OK6>unc-13^RNAi^(attP2*)) or Rbp (*OK6>Rbp^RNAi^(attP2))*. Principal component analysis of normalized libraries (**Figure 2B**) shows that the samples of each genotype cluster together, as expected. However, samples from *OK6>Rbp^RNAi^(attP2)* and *OK6>unc-13^RNAi^(attP2*) are located closer to each other on the first principal component. This represents the axis of greatest variation between the data sets, indicating that the transcriptomes of RSSP knockdowns are more similar to each other than they are to the control.

**Figure 2.**
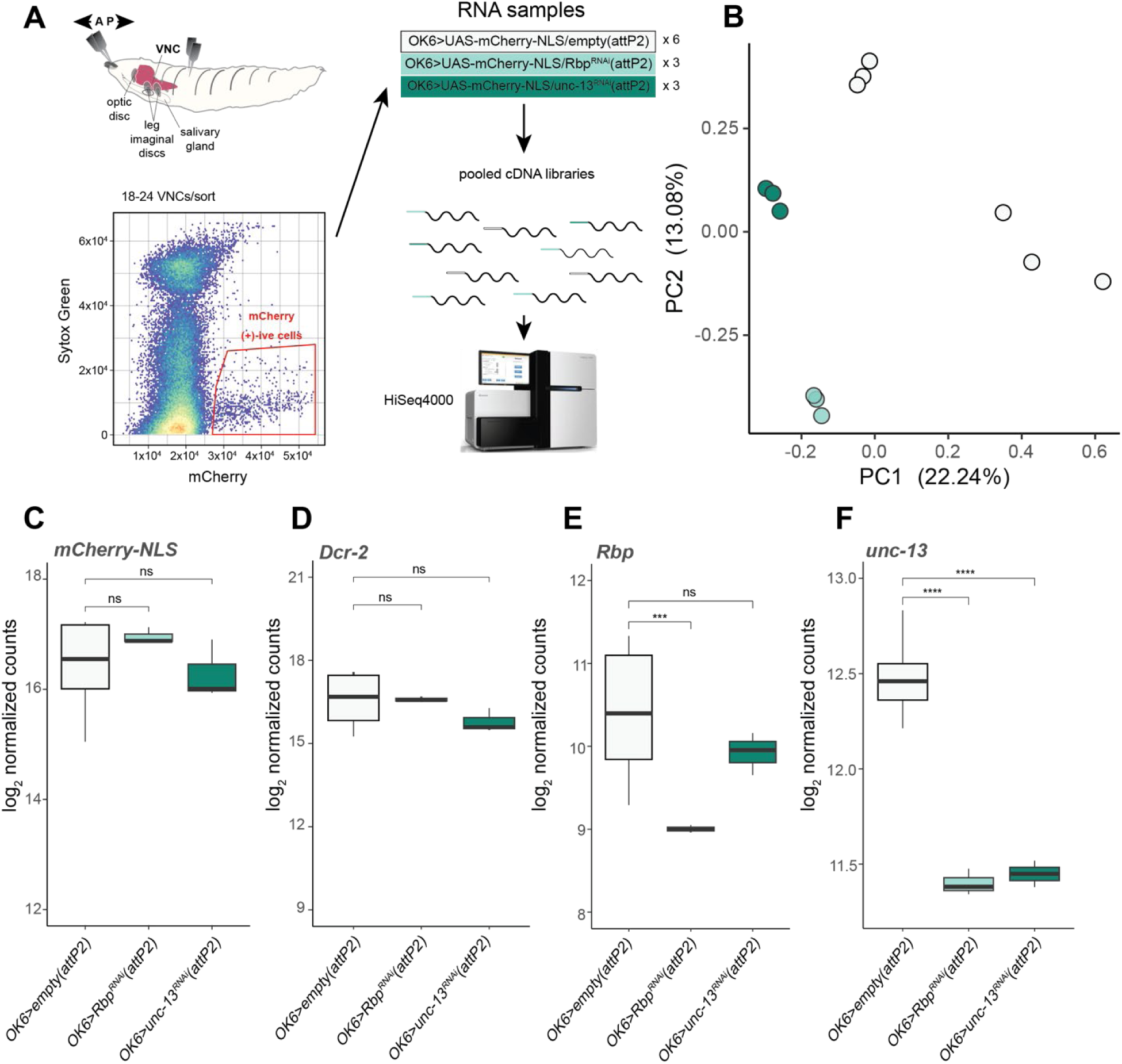
Knockdown RSSPs produce transcriptionally similar phenotypes. **A)** Schematic for generating RNA-seq libraries from sorted motor neuron populations from control (*OK6>empty(attP2)*) and RNAi (*OK6>Rbp^RNAi^(attP2)*, *OK6>unc-13^RNAi^(attP2)*) animals. **B**) Principal component analysis (PCA) plot of RNA-seq samples points colored as depicted in **A**. Axes are annotated by percent variance explained for each principal component. **C-F**) Boxplots of log2 normalized pseudocount expression of **C**) *mCherry-NLS*, **D**) *Dcr-2*, **E**) *Rbp*, and **F**) *unc-13* (BH-adjusted Wald test *p*-values, ns *p* > 0.05, * *p* < 0.01, ** *p* < 0.001, *** *p* < 0.0001, **** *p* < 1e-05).

We assessed the effectiveness of normalization by testing for differential expression of *UAS* transgenes and dsRNA-mediated knockdown as ground truths. While the *OK6>unc-13^RNAi^(attP2)* samples have lower expression of the *UAS-Dcr-2* transgene, samples from all three genotypes have the similar expression of the *UAS-mCherry-NLS* reporter (**Figure 2C, D**). *OK6>Rbp^RNAi^(attP2)* samples had a 2.5-fold decrease in *Rbp* expression compared to the *OK6>empty(attP2)* controls, while the *OK6>unc-13^RNAi^(attP2)* samples had no difference in *Rbp* expression (**Figure 2E**). Interestingly, both the *OK6>unc-13^RNAi^(attP2)* and the *OK6>Rbp^RNAi^(attP2)* had an approximately 2-fold decrease in *unc-13* expression (**Figure 2F**).

Intrigued by the finding that loss of *Rbp* resulted in a concomitant decrease in *unc-13* at the transcriptional level, we sought to quantify expression of Unc-13 at the AZ. To do this we crossed the RNAi and empty *attP2* controls into an *OK6>Gal4;SynapGCaMP6f* line, which bears a GCaMP6f construct localized to the postsynaptic membrane^5^. We labelled larval fillet preparations with an anti-GFP antibody to identify type Ib MN terminals, an anti-Brp antibody to identify AZs and an antibody against Unc-13-A. Since the diameter of a Brp punctum in a wild-type Ib bouton is on the order of a few hundred nanometers^21,22^, we measured the fluorescence intensity by confocal imaging with Airyscan detection for improved spatial resolution (**Methods**). We found that Brp molecules were arranged in small, ring-like clusters as shown previously^21^ (**Figure S2A-C**) and that the Unc-13A intensity at the AZ was reduced by ∼67% in the *Rbp* RNAi and 74% in the *unc-13* RNAi (**Figure S2D**).

### Knockdown of RSSP components increases expression of genes encoding neuropeptides and their secretion machinery

In order to gain insight into the homeostatic alterations following synaptic perturbations, we asked what genes are differentially expressed in each RNAi compared to the *OK6>empty(attP2)* control. Since PC1 accounts for ∼22% of variation in the transcriptomes for each genotype and represents the axis of greatest separation between the controls and RNAi knockdowns (**Figure 2B**), we first examined the genes that contribute to the greatest loadings to this component (**Figure 3A**). Examining the top 50 genes, we found the greatest magnitude of weights encoding neuropeptides (*NPF*, Neuropeptide F; *Nplp2*, Neuropeptide-like precursor 2), secreted proteinaceous ligands (*Swim*, Secreted Wnt-interacting molecule; *Idgf6*, Imaginal disc growth factor 6), ion channels and transporters (*ppk,* pickpocket; *ppk26*, pickpocket 26) and cellular adhesion proteins (*Fit2*, Fermitin 2; *LanA*, Laminin A; *Muc14A*, Mucin 14A). These results suggest that diverse changes in ion transport, excitability and neuromodulatory signaling are common features of weakened excitatory transmitter release.

**Figure 3.**
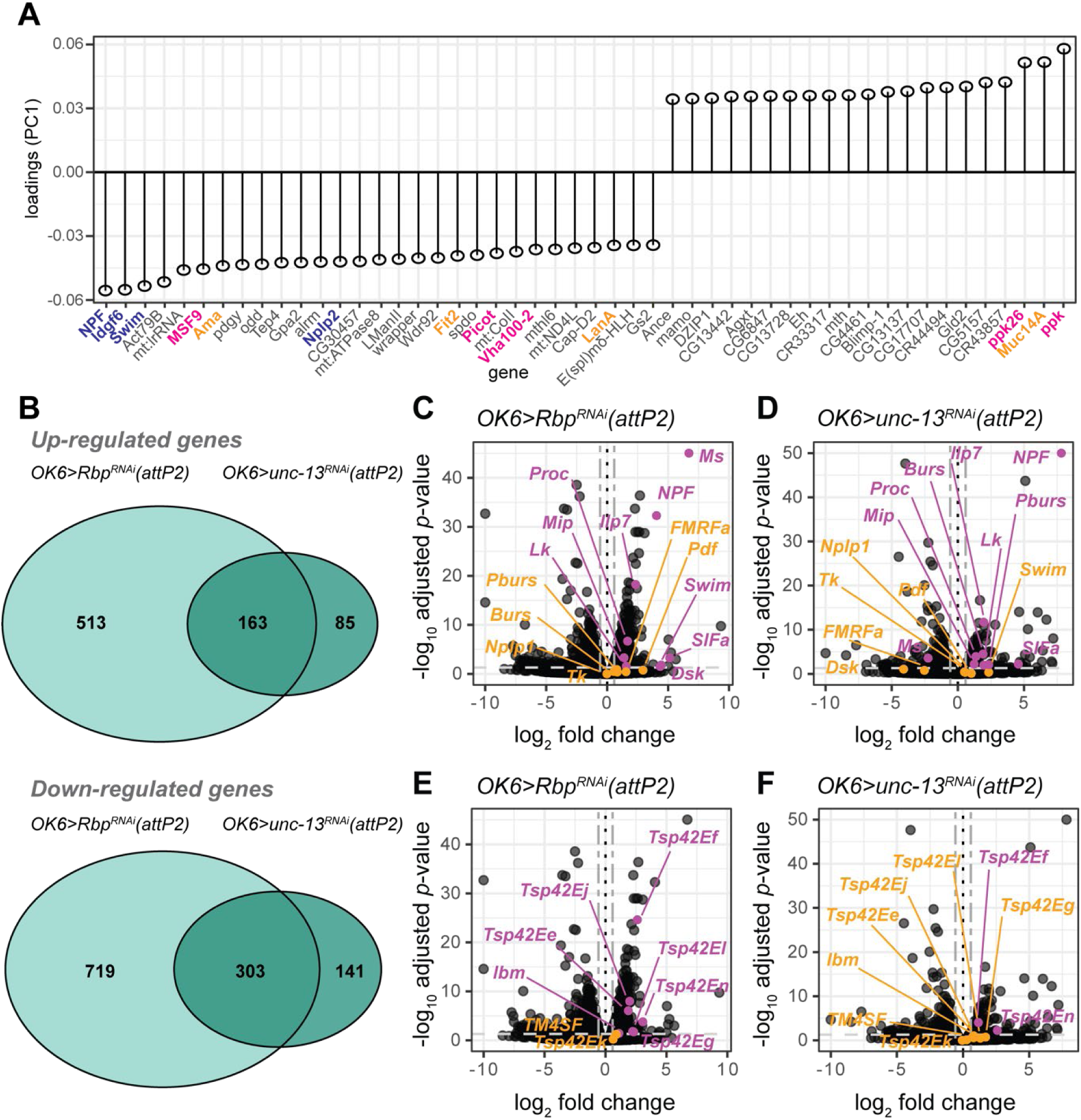
Rbp and unc-13 knockdowns yield overlapping changes in gene expression. **A)** Weights of the top 50 genes contributing to PC1 ranked by magnitude (adjusted p-value < 0.05). Gene names in blue, magenta, and orange are neuropeptides and secreted proteins, ion channels and transporters, and cell-cell or cell-matrix adhesion proteins. **B)** Venn diagram of differentially expressed genes (FDR = 0.05) by knockdown condition. **C, D)** Volcano plots of *OK6>Rbp^RNAi^* () and *OK6>unc-13^RNAi^* **(C)** highlighting genes encoding neuropeptides and secreted proteins**. D, E)** Volcano plots of *OK6>Rbp^RNAi^***(D)** and *OK6>unc-13^RNAi^* **(E)** highlighting genes encoding exosome markers. Black dotted vertical lines in figures B-E demarcate a log_2_-fold-change of zero. The light gray single-dashed horizontal line demarcates an adjusted *p*-value of 0.05. The dark gray double-dashed vertical lines demarcate a fold-change of 0.5. Data points highlighted in magenta have an adjusted *p*-value < 0.05, while points highlighted in orange have a *p-*value ≥ 0.05.

We next performed differential expression analysis to further characterize common features shared by each knockdown. We found that the *OK6>Rbp^RNAi^(attP2)* samples had a greater number (1311 vs 459) of differentially expressed genes compared to *OK6>unc-13^RNAi^(attP2)* samples (**Figure 3A**). However, the majority of differentially expressed genes identified in the *OK6>unc-13^RNAi^(attP2)* samples were also found to be differentially expressed in the *OK6>Rbp^RNAi^(attP2)* (**Figure 3B**), suggesting that *Rbp* has an upstream genetic interaction with *unc-13*. Examining this set of genes with increased expression compared to *OK6>empty(attP2)* controls, we found that genes encoding an overlapping set of neuropeptides were up-regulated in both *OK6>Rbp^RNAi^(attP2)* and *OK6>unc-13^RNAi^(attP2)* (**Figure 3C,D; Supplementary Tables 2-3**). The biggest increases in expression, which were shared between the two knockdowns, were for NPF and SIFa, whose central actions regulate a wide range of behaviors, including circuits controlling sleep, learning and memory, feeding and metabolic homeostasis^23^. Proctolin, which is found in a subset of type Ib MNs and their presynaptic nerve terminals, as well as in the type Is MNs projecting to dorsal muscle segments also increased in expression^24,25^, but to a lesser extent. We detected receptors in the type I MNs for two of these neuropeptides, NPFR and SIFaR, but not for proctolin (**Supplementary Tables 4-5**), which has been shown to act postsynaptically^25^.

Since neuropeptides and other protein ligands are secreted from *Drosophila* motor neurons from large dense core vesicles or exosomes^26,27^, we also examined genes in the tetraspanin family, which are broadly involved in membrane organization and trafficking, but have specific roles in modulating glutamatergic transmission in the mammalian CNS^28^. We found that several of the tetraspanins were up-regulated in *OK6>Rbp^RNAi^(attP2)* (**Figure 3E; Supplementary Table 2**) and that two of those, *Tsp42En* and *Tsp42Ef*, were also up-regulated in *OK6>unc-13^RNAi^(attP2)* (**Figure 2F**). We confirmed the increase in expression of *Tsp42Ef* with RT-PCR (**Figure S3A**).

### Knockdown of RSSP components decreases expression of AZ structural components

Our finding that the *Rbp* and *unc-13* knockdowns change the expression of a genes involved in peptidergic signaling and trafficking led us to wonder whether altered expression of genes encoding other components of the transmitter release machinery could contribute to weakened transmitter release. In both knockdowns, we observed decreased expression of *cac,* as well as of the cytomatrix AZ protein *Rim* (**Figure 4**) ^15–18^. The *Rbp* knockdown had altered expression of additional genes encoding proteins that modulate the coupling between calcium entry and synaptic release. These genes include decreased expression of *unc-13*, as already mentioned, and of *straightjacket* (*stj*), the α2δ auxiliary subunit of the Cac channel^29^, as well as increased expression of the fusion clamp *complexin* (*cpx*)^30^, *Ras opposite* (*rop*)^31^, the homolog of mouse *Unc18*)^32^, *αSnap*^33^, *comatose* (*comt*), the homolog of mouse *Nsf*^34^, and the A-kinase *Pka-C1*^35,36^ (**Figure 4**). Since Rbp, Unc-13, and RIM regulate coupling between synaptic vesicles and Cac, and Stj stabilizes Cac, these proteins collectively form the keystone of AZ organization^29,37^. Their decrease in expression, along with the increase in the Cpx fusion clamp, are consistent with the observed decrease in neurotransmitter release^38^ (**Figure 1**). These results were confirmed by RT-PCR for both *cac* and *Rim* (**Figure S3B**) and by antibody staining followed by Airyscan confocal imaging for Cac (**Figure S4**).

**Figure 4.**
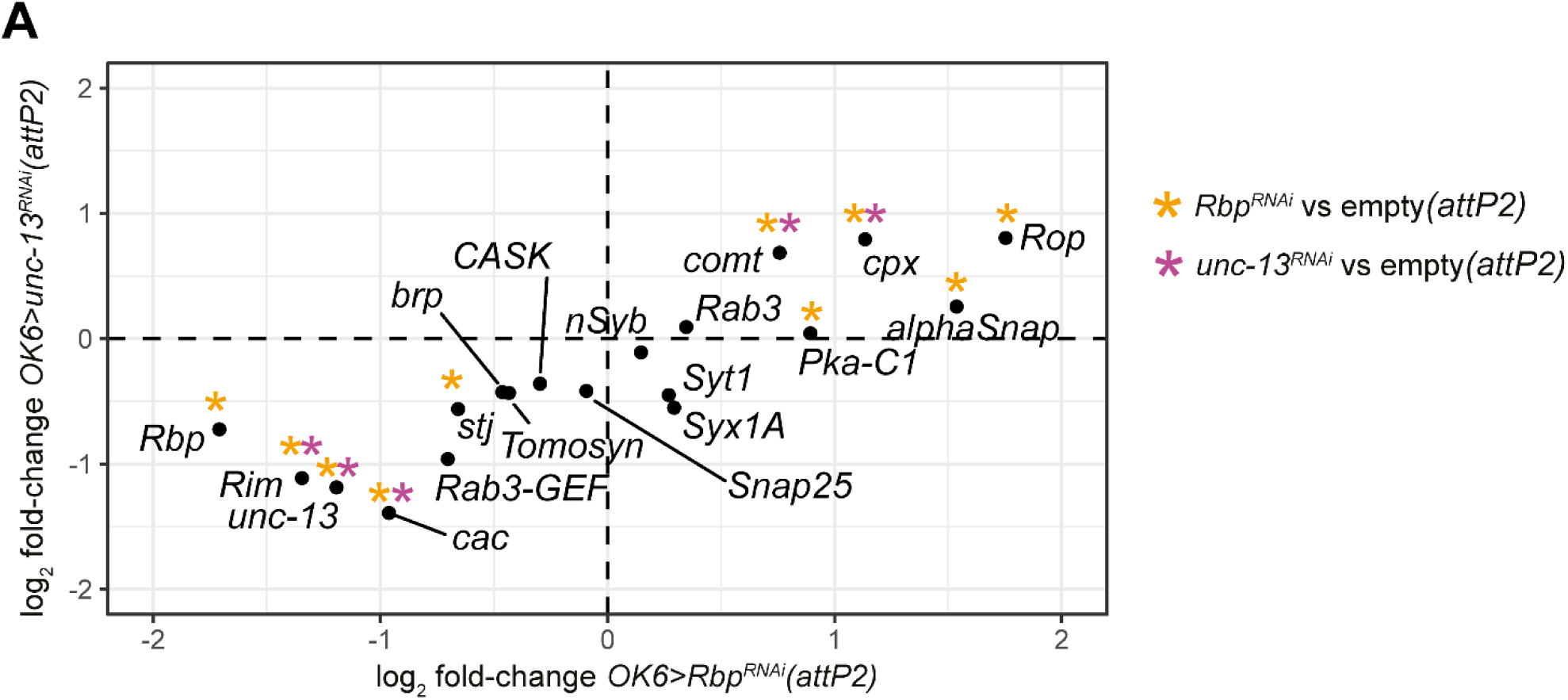
Knockdown of RSSPs decreases the expression of AZ components. **A)** log_2_ fold-change of *OK6>Rbp^RNAi^(attP2)* (horizontal axis) and *OK6>unc-13^RNAi^(attP2)* (vertical axis) highlighting genes comprising presynaptic transmitter release machinery. The log_2_ fold-change in expression for each genotype is compared to the *OK6>empty(attP2)* control. Black dotted lines demarcate a log_2_-fold-change of zero. * indicates an adjusted *p-*value < 0.05. Asterisks in amber are DE in *OK6>Rbp^RNAi^(attP2)* and asterisks in plum are DE in *OK6>unc-13^RNAi^(attP2)*.

### Knockdown of RSSP components decreases expression of genes encoding Kv potassium channels and auxiliary subunits

In effort to understand how perturbations that reduce glutamate release by MNs increase their firing^4,10^, we asked if the *Rbp* and *unc-13* knockdowns also have altered expression of genes that regulate excitability. We found that, in both the *Rbp* and *unc-13* knockdowns, MNs had reduced expression of three voltage-gated (K_v_) K^+^ channels, whose reduced function increases excitability: the A-type *Shaker* (*Sh*, K_v_1) channel^39^, the delayed-rectifier *Shab* (K_v_2) channel^40^ and the *ether a go-go* (*eag*) family member *Elk* channel^41^ (**Figure 5, Supplementary Tables 6-7**). The KCNQ (K_v_7) channel also had reduced expression, but only in the *Rbp* knockdown (**Figure 5, Supplementary Table 6**). There was no change in expression of three other K_v_ channels: *Shal* (K_v_3), *Shaw* (K_v_4) and *eag* itself. Expression was also reduced in *Slob*, an auxiliary subunit of the *Slo* (BK) voltage and Ca^2+^ activated K^+^ channel^42^, but this only occurred in the *Rbp* knockdown (**Figure 5, Supplementary Table 6**).

**Figure 5.**
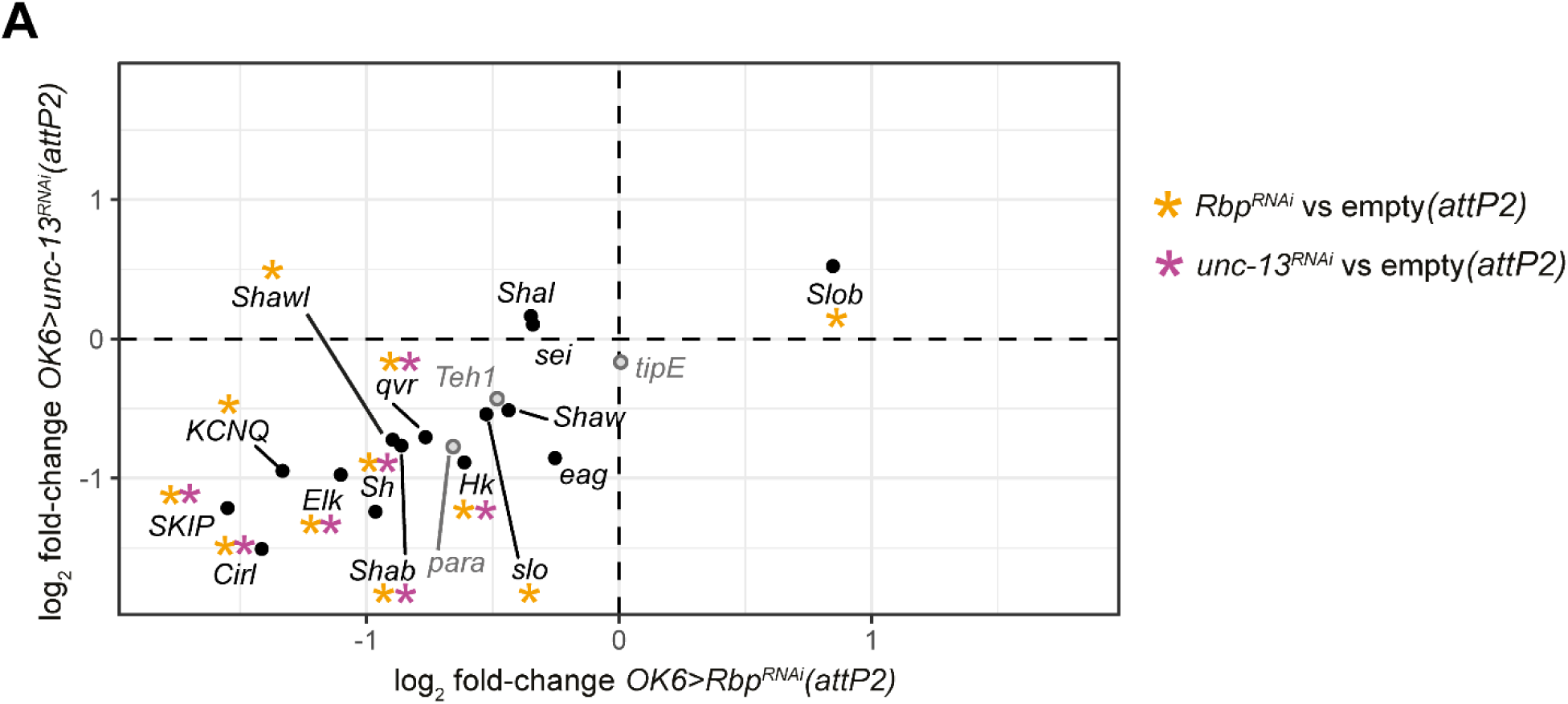
Knockdown of RSSPs results in down-regulation of suppressors of excitability. **A)** log_2_ fold-change of *OK6>Rbp^RNAi^(attP2)* (horizontal axis) and *OK6>unc-13^RNAi^(attP2)* (vertical axis) highlighting genes encoding Kv channels and the aGPCR, Cirl (black) and Nav channels and associated proteins (grey). The log_2_ fold-change in expression for each genotype is compared to the *OK6>empty(attP2)* control. Black dotted lines demarcate a log_2_-fold-change of zero. * indicates an adjusted *p-*value < 0.05. Asterisks in amber are DE in *OK6>Rbp^RNAi^(attP2)* and asterisks in plum are DE in *OK6>unc-13^RNAi^(attP2)*.

We next examined the expression of channel regulators and found that both the *unc-13* and *Rbp* knockdowns had decreased expression of three positive regulators of K_v_ channels: *Hyperkinetic* (*Hk*), *quiver* (*qvr/SSS) and SKIP* (**Figure 5, Supplementary Tables 5-6**), whose loss reduces K+ current and increases neuronal excitability. In addition to these changes in genes encoding K+ channels and their auxiliary subunits, both the *unc-13* and *Rbp* knockdowns had a reduction in the expression of Cirl (Ca^2+^ Independent Receptor for Latrotoxin) (**Figure 5, Supplementary Table 6**), a Gi-coupled adhesion GPCR whose loss may contribute to increased firing^43^.

In contrast to the decreased expression of K^+^ channels and their modulators in the RSSP knockdowns, there was no change in expression of the voltage-gated sodium channel (*para*)^44^, or of two *para* β subunits, *tipE*^45,46^ and *Teh1*^47^, or of a *para* genetic regulator, *pasilla* (*ps*)^48,49^ (**Figure 5, Supplementary Tables 5-6**). We confirmed the decreased expression of *SKIP*, *Shab*, *qvr* and *Hk* by RT-PCR, using *eag* as a control (**Figure S3C**).

We also asked if RSSP knockdown results in altered expression of ionotropic neurotransmitter receptors that could affect the response to synaptic input from pre-motor interneurons. *Drosophila* interneurons fall into three classes—cholinergic (excitatory), GABAergic (inhibitory) or glutamatergic (primarily inhibitory via *GluCl*α glutamate-activated Cl-channels)^50–55^. We examined the expression genes encoding synaptic receptors to glutamate, acetylcholine or GABA, which represent the major transmitters that pre-motor inputs release onto MNs^56^. We found no change in expression for the majority of genes encoding transmitter receptors (**Supplementary Tables 10-11**). However, in both the *unc-13* and *Rbp* knockdowns, we observed decreased expression of the excitatory AMPA-type ionotropic glutamate receptor *GluRIB* and of the excitatory nicotinic acetylcholine receptor *nAChRalpha6*, but also of the inhibitory GABA-A receptor subunit *Rdl* (**Supplementary Tables 10-11**). Moreover, in addition to its effect on *Shaker* current, *qvr/SSS* inhibits acetylcholine receptor activity^57^, so that reduced *qvr/SSS* expression increases cholinergic synaptic excitation. The result of these changes is mixed: two changes that favor synaptic excitation over inhibition and two changes that have the opposite effect, leading to no clear outcome. Thus, we observe an orchestrated decrease in expression in type I MNs of three K_v_ channels and five positive modulators of K_v_ channels.

## Discussion

### Homeostatic coupling of AP firing to synaptic transmission

Following changes to neural activity or synaptic transmission that occur during development or as a result of synaptic plasticity, neuromodulation or pathological perturbation, homeostatic mechanisms adjust neural properties to return AP firing and synaptic transmission to set points that ensure proper circuit output^1–3^. At the model glutamatergic synapse of the *Drosophila* larval NMJ, retrograde signaling from postsynaptic muscles to presynaptic type I MNs triggers PHP, an increase in AP-evoked glutamate release that compensates for a decrease in the amplitude of the excitatory postsynaptic potential due to either a null mutation of the GluRIIA subunit of the GluRII ionotropic glutamate receptor or to pharmacological block of GluRII^3,7^. However, the boost in glutamate release only occurs in only one of the two convergent type I glutamatergic MN inputs to body wall muscle, the type Ib MN, and not in the type Is MN, which elicits more than half of the amplitude of the combined EPSP^8,9^. In other words, this PHP mechanism only partially compensates for weakened synaptic transmission^21^. Despite this, locomotory behavior of the *GluRIIA^-/-^*animals remains near normal^11^. Moreover, animals also locomote near normally when they have presynaptic knockdown of any one of three components of the glutamate release machinery—*unc-13, Rbp* or *cac*—which severely reduces AP-evoked release, without detectible postsynaptic compensation^11^. These presynaptically and postsynaptically weakened animals show a common change in MN activity pattern: they increase the duration of activity bursts that drive the locomotory peristaltic waves of contraction. This indicates that there is a second mechanism of homeostatic regulation that compensation for weakened per AP synaptic transmission from MN to muscle in which the MNs fire more APs, as suggested earlier^7^.

We probed the mechanism of MN firing homeostasis by characterizing the transcriptional consequences of impaired glutamate release due to knockdown of the same presynaptic components of the transmitter release machinery, *unc-13* and *Rbp*. We find that knockdown of *unc-13* or *Rbp* changes the expression of an overlapping set of genes, and knockdown of *Rbp* itself reduces *unc-13* expression, suggesting that there is a hierarchical relationship between the two RSSPs and that both the overlapping changes in gene expression and much of the presynaptic weakening are due to the reduction in *unc-13.* Among the genes that changed in expression level in the two RSSP knockdowns that weakened transmitter release from type I MNs, two sets of genes stood out: genes encoding K_v_ channels and their positive modulators and genes encoding neuropeptides and proteins implicated in neuropeptide release.

### Changes in expression of genes encoding channels and modulators that regulate excitability

Knockdown of *unc-13* or *Rbp* decreased the expression of a group of genes that encode pore forming α-subunits of K_v_ channels or modulatory proteins that enhance their function. These include: the inactivating (I_A_) Shaker (K_v_1) channel, the delayed rectifier Shab (K_v_2) channel and the Eag-like Elk (K_v_12) channel. Additionally, in the *Rbp* knockdown, we also see reduction in expression of the voltage- and Ca^2+^-gated Slo (BK) and voltage-gated Shawl (K_v_4) and KCNQ (K_v_7) K^+^ channels. Reduced K^+^ channel expression is known to increase the excitability of type I MNs^58,59^. More important still, Shaker and Shab mutants have each been shown not only to increase AP firing, but to increase the duration of MN locomotory bursts^60^, exactly as observed in the intact larva during restrained locomotion with knockdown of *unc-13* and *Rbp*^11^. In other words, reduced expression of these K_v_ channels can account for the compensatory alteration of firing patterns that support locomotor behavior.

Both the *unc-13* and *Rbp* knockdowns also reduced expression of three genes encoding K^+^ channel modulatory subunits that boost K_v_ channel function: the Shaker interacting proteins *Hk* and the Shaker and *qvr/SSS,* as well as the Shal regulator SKIP. *Hk* is a β subunit of the Shaker channel, which boosts channel current amplitude, accelerates current rise and shifts the voltage dependence of activation to more negative voltage^61,62^. Elimination of *Hk* resembles reduction of *Shaker* expression and leads to hyper-excitability. *Qvr/SSS* is an Ly6/GPI-anchored protein, whose loss slows Shaker channel current rise and increases cumulative inactivation, resulting in a decreased Shaker current amplitude and increased excitability^64^. *SKIP* slows the inactivation of Shal channels and its elimination results in fast inactivation (*i.e.* reduction of K+ current) and greater excitability^65^. Expression was also reduced in *Slob*, an auxiliary subunit of the *Slo* (BK) voltage and Ca^2+^ activated K^+^ channel^42^, but this only occurred in the *Rbp* knockdown (**Figure 5, Supplementary Table 6**). Deletion or knockdown of *Slob* increases AP firing in Drosophila type I MNs^66^ and enhances glutamate release^67^. In addition, both the *unc-13* and *rbp* knockdowns had a reduction in the expression of Cirl (Ca^2+^ Independent Receptor for Latrotoxin) (**Figure 5, Supplementary Table 6**). Cirl is a Gi-coupled adhesion GPCR whose activity reduces cAMP levels and protein kinase A activity in cells^68^. Loss of function of Cirl may also contribute to increased firing^43^.

It should be noted that we did not observe complete loss of expression of these genes, only reductions by 30-65% compared to controls. However, increased excitability is also seen in hypomorphs and heterozygotes with one wildtype copy of the gene^69,70^. This suggests that reduction by ∼50%, as we observe here in the *unc13* and *Rbp* knockdowns, is sufficient to increase firing, even when it takes place in only one of the K_v_ module genes. Since we observe a simultaneous decrease of this magnitude in multiple genes, the increase in excitability is expected to be substantial.

Taken together, weakening of glutamate release by type I MNs coordinately reduces expression of genes encoding K_v_ channels and their function-boosting associated proteins. We propose that co-suppression of this gene module functions as a mechanism for increasing the number of APs in locomotory bursts to compensate for reduced glutamate release per AP. Intriguingly, three of these channels—Shaker, Shab and Slo—have already been found to coordinately increase in expression in response to elimination of the Shal (K_v_3) K^+^ channel^71^, supporting the notion that they are part of a co-regulated transcriptional module.

### Changes in expression of neuropeptide genes

A second set of genes that showed large changes in expression and similar biological function caught our attention in the RSSP knockdowns. These encode neuropeptides and tetraspanins, which are implicated in neuropeptide exocytosis^26,27^. All of these genes had increased expression in the RSSP knockdowns. Both *Drosophila* and mammalian neurons commonly release small molecule neurotransmitters from small clear vesicles, with a reasonable probability of release at both low and high frequencies of AP firing, and neuropeptides from large dense core vesicles during high frequency AP trains^23,72^. The neuropeptides can act pre- or postsynaptically and expand the dynamic range of the synapse^73,74^. Single-cell transcriptomic atlases of the adult fly brain have shown co-expression of *VGlut*, indicating glutamatergic neurons, with the genes encoding neuropeptides *Nplp1*, *Mip*, *Dh44*, *Proct*, *CCHa1*, *CCHa2* and *FRMFa*^75^.

In the RSSP knockdowns, we observed increased expression of the genes encoding Neuropeptide F (NPF), the *Drosophila* homolog of neuropeptide Y, SIFa and Proloctin (Proc). NPF and SIFa, which regulate a wide range of behaviors^23^, increased expression by 4 to 7-fold, enough to place them into the top 20 differentially expressed genes. However, a recent study^76^ showed that they are both expressed at very low levels, suggesting that these changes may not biologically meaningful. In contrast, Proctolin, which is expressed at high levels^76^ has been shown to increase contractions of body wall muscles^77,25^. This postsynaptic enhancement of contraction could help to compensate for weakened synaptic transmission to help preserve locomotion.

## Conclusion

Weakening of glutamatergic transmission from *Drosophila* MNs to muscle engages two multiplicative mechanisms of homeostasis: PHP, which boosts the amount of transmitter release per AP^4^, and increased duration of MN bursts, which can further boost release by increasing the number of APs that drive the contraction wave of locomotion^10,11^. We find here that molecular disruptions that reduce AP-evoked glutamate release drive a coordinated reduction in expression of a group of K_v_ channels and modulatory proteins that augment K_v_ channel function. These transcriptional changes provide a molecular mechanism for the compensatory increase in MN locomotory burst duration and can work multiplicatively with the PHP increase in release per AP by increasing the number of APs, thereby preserving locomotion in the face of synaptic perturbations.

## Methods

### Fly Stocks

The *UAS-Rbp^RNAi^* line (TRiP.JF02471, BDSC #29331), *UAS-unc-13^RNAi^* line (TRiP.JF02440, BDSC #29548), *UAS-cac^RNAi^* line (TRiP.JF02572, BDSC #27244) and the empty *attP2* acceptor line (BDSC #36303) were obtained from the Bloomington Drosophila Stock Center as donated by the Transgenic RNAi Project. *UAS-dsRNA* constructs were driven in type I MNs with *OK6-Gal4*^13,79^. *UAS-Dcr2* (BDSC #24646) was used to improve the efficacy of RNA interference^80^. Fruit fly stocks bearing a synaptically-localized GCaMP6f (*SynapGCaMP*) under the control of the myosin heavy-chain (*MHC*) promoter were used for reporting postsynaptic activity in the muscle^5^ in electrophysiology experiments. For RNA-seq experiments, fruit fly stocks bearing transgenes encoding nuclear mCherry (*UAS-mCherry-NLS*, BDSC #38424, ref ^81^), along with Dcr2 and dsRNA constructs, in type I MNs were used for the purpose of FACS sorting.

Larvae were raised to the 3rd instar stage on cornmeal and molasses media in an incubator at 25°C. Female parents bearing the *OK6-Gal4* driver and *UAS-Dcr2* transgenes were crossed with male parents bearing either the *attP2* empty vector or RNAi construct to obtain control and knockdown progeny, respectively. 3rd instar larva with the following genotypes were used for RNA-seq: *UAS-Dcr2;OK6-Gal4/+;UAS-mCherry-NLS/(attP2)* (*abbr. OK6>empty(attP2)*) *UAS-Dcr2;OK6-Gal4/+;UAS-mCherry-NLS/UAS-Rbp^RNAi^(attP2)* (abbr. *OK6>Rbp^RNAi^(attP2)*) *UAS-Dcr2;OK6-Gal4/+;UAS-mCherry-NLS/UAS-unc-13^RNAi^(attP2)* (abbr. *OK6>unc-13^RNAi^(attP2)*)

3^rd^ instar larva of the following genotypes were used for immunohistochemistry and optical quantal analysis experiments:

*UAS-Dcr2;OK6-Gal4/+;MHC-SynapGCaMP6f/(attP2)*

*UAS-Dcr2;OK6-Gal4/+;MHC-SynapGCaMP6f/UAS-Rbp^RNAi^(attP2)*

*UAS-Dcr2;OK6-Gal4/+;MHC-SynapGCaMP6f/UAS-unc-13^RNAi^(attP2)*

*UAS-Dcr2;OK6-Gal4/+;MHC-SynapGCaMP6f/UAS-cac-13^RNAi^(attP2)*

SynapGCaMP6f Optical quantal imaging

### Optical quantal imaging

Optical quantal imaging was performed similarly to our previous reports^19,20^. To summarize, third instar larvae were dissected on PDMS (Sylgard 184, Dow Corning, Auburn, MI) pads in ice-cold HL3 solution containing, in mM: 70 NaCl, 5 KCl, 0.45 CaCl2, 20 MgCl2, 10 NaHCO3, 5 trehalose, 115 sucrose, 5 HEPES, and with pH adjusted to 7.2. Following removal of the brain (VNC), larval fillets were washed and imaged in room temperature HL3 containing 1.5mM Ca^2+^ and 25mM Mg^2+^. Fluorescence images were acquired at room temperature with a Vivo Spinning Disk Confocal microscope (3i Intelligent Imaging Innovations, Denver, CO), using a 63 × 1.0NA water immersion objective (Zeiss), 1.2X optical adapter, LaserStack 488 nm (50 mW) laser, CSU-X1 A1 spinning disk (Yokogawa Tokyo, Japan), standard GFP filter, and EMCCD camera (Photometrics Evolve512, Tucson, AZ). All live SynapGCaMP6f imaging recordings were done on ventral longitudinal abdominal muscle 4 at segments A3-A5 of third instar larvae. All imaging was performed using 50 ms exposures (20 fps) of the full camera sensor (512 × 512 px). Nerve stimulation was performed with a suction electrode attached to a Stimulus Isolation Unit (SIU, ISO-Flex, A.M.P.I. Jerusalem, Israel), with 100 us stimulus duration. The intensity of the stimulus was adjusted to recruit both Ib and Is axons (verified through imaging) and kept constant throughout the imaging session. Nerve stimulation and imaging were synchronized using custom-written Matlab scripts (Matlab Version 2015b, MathWorks, Inc., Natick, MA) in order to control the SIU and trigger imaging episodes with SlideBook (v6.0.16, 3i Intelligent Imaging Innovations). For comparability between experiments, recordings were done on only one NMJ (Ib-Is pair) per larva and recordings were performed within 30 minutes from the start of the dissection, ensuring the animals health. As in our previous report^20^, we alternated which segments (A3-A5) we imaged from since no significant difference was found between segments.

### Functional registration and bleach correction

The initial quantal image analysis was performed using custom-written MATLAB protocols, same as in our previous work^20^. Individual stimulus episodes were excluded due to out of focus NMJs, moving NMJs or failed axon recruitment. Otherwise, all movies were filtered (Gaussian low-pass filter), to reduce high-frequency noise. Image analysis areas were then separated into Ib and Is NMJ regions. All imaging data were registered using a multi-stage approach, during which all images were registered to a common reference image (usually the first frame of the first image).

Following area selection and reference image selection, NMJs were tracked relative to this reference image using a rigid subpixel registration method to remove any large movements within the NMJ imaging area^19,20,35^. As in our previous work^20^, we corrected for local bouton movements using a custom diffeomorphic implementation of a demons algorirthm^36^ Once motion corrected, movies were bleach corrected using a fit for a double exponential bleach correction curve to the mean baseline pre- and post-stimulus fluorescence data for each trial separately. Following bleach correction, both ΔF and ΔF/F movies were generated using the first image as the baseline fluorescence (F0) image for the episodic data.

### Quantal synaptic optical reconstruction (QuaSOR)

After event detection, isolation, and verification, we then proceeded to analyze the 2D response profile of each event’s maximum ΔF/F spatial profile using the custom QuaSOR algorithm as in our previous work^20^. Briefly, we first looked at all the identified responses from our quantal detection and isolated small ROIs containing individual or small groups of partially overlapping response fields that corresponded to individual or small groups of events. These smaller ROIs were then subjected to independent 2D Gaussian mixture model fitting of the isolated ΔF/F spatial profiles. Gaussian mixture model fitting for all response ROIs, all event functions were then remapped onto a common coordinate space and merged to define a single set of 2D Gaussian functions for each quantal response. The peak positions of each 2D Gaussian component were used to define event locations in a 21.2 nm x 21.2 nm pixel coordinate space. For visualization purposes, maps were generated by applying a normalized 2D Gaussian filter to each event coordinate prior to adding each event to the overall image. In this way each pixel contains an approximation of the event density at that location. Local QuaSOR synapse alignments were performed by identifying maximum evoked coordinate density positions for synaptic ROIs within a 350 nm radius. Maintaining their relative organization to nearby events, these QuaSOR event coordinates were averaged together with other synapses to generate a mean density image for synapse groupings.

### CNS Dissection

Prior to dissection 3rd instar larva were screened for the presence of nuclear mCherry in the VNC with ZEISS Axio Zoom v16 FL epifluorescence microscope. For bulk RNA-seq experiments, larvae were rinsed twice in 1X PBS (Gibco), once in 70% ethanol, and again in 1X PBS to sterilize the surface of the animals. Larvae were dissected on a Sylgard pad in a droplet of Schneider’s complete media (Gibco) supplemented with 10% FBS (Corning) and 1% PenStrep (Gibco). After removing extraneous tissues, the brain and VNC were transferred to a low-binding 1.5 mL tube containing Schneider’s complete media on ice until dissociation. Dissected tissues were kept on ice prior to dissociation no longer than 60 minutes to ensure the viability of the cells during FACS sorting.

### CNS Dissociation

Tissue samples were washed three times with 500 μL Schneider’s media containing 1.5 mM EDTA and 1.5 mM L-Cysteine to remove media containing FBS. Fresh dissociation solution was prepared for each experiment from Schneider’s media with the following additives: 1.5 mM EDTA, 1.5 mM L-Cysteine, papain (Worthington), and 1.25 mg/mL collagenase type-I (MilliporeSigma). Tissue samples were transferred into the dissociation solution and incubated at 25°C on a nutator for 5 minutes. Samples were triturated 30 times with a P200 pipette tip. This process of incubation and trituration was repeated before a final step in which tissue was sheared 7 times with a 27G1/2 needle.

The digestion reaction was quenched with the addition of 500 μL Schneider’s complete media with 10% FBS and 1% PenStrep. Dissociated cells were passed through a 40 μm cell strainer (Falcon) and transferred to a low-binding 1.5 mL tube. The processed sample was centrifuged at 0.7 x *g* for 7 minutes, after which the supernatant was replaced with 1 mL of Schneider’s complete media containing 5% FBS and 1% PenStrep.

### Fluorescence Activated Cell Sorting

Processed cell samples were stained with 0.3 μM Sytox Green (Molecular Probes) to assay cell viability prior to sorting. Cells were sorted on a BD Influx Cell Sorter with 140 μm tubing at pressures < 12 PSI. The sorted population was chosen for high expression of nuclear mCherry (PE-Texas Red) and low Sytox Green fluorescence (GFP). Plots of FACS data were created using the flowCore (ver 1.52.1)^82^ and ggcyto packages (ver 1.14.0)^83^.

### Low-input RNA-seq Library Preparation and Sequencing

Cells were sorted into RLT lysis buffer (RNeasy Micro, QIAGEN) supplemented with 3.5 mM BME. Cells were lysed by vortexing for 30 seconds at room temperature. RNA extraction was performed according to RNEasy Micro kit protocol, omitting the DNase digestion step. Extracted RNA was checked for quality by an Agilent 2100 Expert Bioanalyzer with a Eukaryote Total RNA Pico chip.

Sequencing libraries were prepared with the SMART-Seq2 protocol with modifications to RNA capture and cDNA amplification steps. RNA was annealed to an oligo-dT primer (5’-AAGCAGTGGTATCAACGCAGAGTACT_30_VN-3’, IDT) in a buffer containing 2.5 mM dNTPs and recombinant RNase Inhibitor (Takara). First-strand synthesis was performed with the SuperScriptII kit using a custom template-switching oligo (5’-/5Me-isodC//iisodG//iMe-isodC/AAGCAGTGGTATCAACGCAGAGTACATrGrGrG-3’, IDT) with 5’ modifications to avoid the creation of concatemers in low input samples^84^. cDNA was digested with lambda exonuclease (New England Biolabs) digestion prior to amplification as described previously^85^. Libraries were constructed from amplified cDNA using the Nextera XT kit (Illumina) and size-selected with AMPure XP beads. Fragments above 700 bps were excluded with Pippin Prep (SAGE Biosciences). Indexed libraries were sequenced on Illumina Hi-Seq 4000 sequencers to produce 100 nt paired-end reads.

### RNA-seq Alignment and Pre-processing

Demultiplexed .fastq files were first analyzed with FastQC (ver 0.11.7)^86^ to check for quality and trimmed with Trimmomatic (ver 0.38)^87^. Trimmed reads were aligned to a custom reference genome built from Berkeley Drosophila Genome Project (dm6, release 98) genome assembly and the pUAST-mCherry-NLS vector sequence for the detection of the mCherry mRNA reads. Samples bearing dsRNA constructs were mapped to custom genomes containing the aforementioned transgenes and the sequence of the dsRNA construct. Mapping was performed with HISAT2 (ver 2.1.0)^88^ and the counts matrix was generated with the featureCounts function from the Subread package (ver 1.6.4)^89^.

### Gene Filtering, Normalization and Differential Expression Analysis

Biological analysis was performed in after using a custom script to import the featureCounts output into a SummarizedExperiment object (ver 1.16.1)^90^. Genes with more than 5 counts in at least 3 libraries were considered detectable. Genes that did not meet this threshold were removed from downstream analysis. Libraries were normalized by calculating a size factors for each sample from the median ratio of gene expression relative to the geometric mean for each gene as implemented in DESeq2 (ver 1.26.0)^91^. Identification of differentially expressed (DE) genes between each knockdown condition was performed with the DESeq2. Visualization of DE analysis was performed with custom scripts built upon ggplot2 (ver 3.2.1)^92^.

### qRT-PCR

FACS and RNA purification sorted cells for qRT-PCR experiments were prepared as described for the low-input RNA-seq experiments. cDNA was reverse transcribed from purified RNA with the SuperScript III First-strand Synthesis System (Invitrogen) using 5 μM oligo(dT)_20_ as a primer to improve selectivity for mRNA in the samples. qPCR was performed with the Power SYBR Green PCR Master Mix (Applied Biosystems) on a BioRad CFX96 Touch Real-Time PCR Detection System. Each reaction was performed in triplicate along with no-template controls.

### Primer Design

Primers for the qPCR experiments were designed using FlyPrimerBank^93^, with the exception of the primer pair targeting EF1a which was previously published in Ponton et al. Primers were selected by their annealing temperature, favoring primer pairs that span regions of the target gene present in all known isoforms and span exonic regions to reduce the likelihood of genomic DNA amplification. Sequences for each primer used in this study are in Supplementary Methods Table 1.

### Quantification of expression

Expression for each gene of interest was determined with the relative quantification method^95,96^ with correction for differences in primer efficiency^96^. aTub84B and EF1a were chosen as reference genes for relative quantification of gene expression across controls and knockdowns.

### Immunohistochemistry

Larval fillets were prepared as described previously^21^. Larvae were fixed at room temperature in Bouin’s fixative (Ricca Chemical Company, Arlington, TX) for 5 minutes and rinsed with ice cold PBS with 0.1% Triton X100 (PBT) to remove trace fixative. Larval fillets were blocked in PBS with 0.1% Triton X100, 2.5% normal goat serum, and 0.02% sodium azide (PBN) for 30 minutes. All antibody incubations were performed in PBN. For primary antibody staining, mouse anti-Brp (nc82; Developmental Studies Hybridoma Bank, Iowa City, IA) was used at 1:150, while rabbit anti-unc-13A^17^ and rabbit anti-Cac^98^ were used at 1:1500. All primary antibody incubations were performed overnight at 4C. Samples were washed three times for 15-20 minutes in PBT at room temperature prior to blocking and secondary antibody labelling. Alexa Fluor Plus 555 goat anti-rabbit (Invitrogen A32732), Alexa Fluor 647 goat anti-mouse (Thermo Fisher A32733) and goat FITC-conjugated anti-GFP (Rockland Immunochemicals, 600-102-215) were used at 1:1500 and incubated with samples at room temperature for 2-3 hours. All larval fillet preparations were washed in PBT as described above and mounted in Vectashield (H-1000; Vector Laboratories, Burlingame, CA) or Vectashield Hardset (H-1400) prior to imaging.

To reduce background fluorescence and amplify signal, modifications to the staining protocol were made for the Cac immunohistochemistry experiments. Larval fillet preparations were incubated in rabbit anti-Cac as described above, followed by a 9-hour wash in PBT at 4C. Samples were then incubated with 1:750 goat, anti-rabbit Biotin F(ab’)2 (Jackson 111-066-144) along with 1:150 mouse anti-Brp as described for the unc-13A immunohistochemistry experiments. Fillet preparations were then washed in PBT as described above, followed by two 1-hour washes in PBT. Samples were blocked and stained with 1:750 Streptavidin-555+ (Thermo Fisher S32355) and 1:1500 donkey anti-mouse 647 (Jackson 715-605-151).

### Airyscan imaging

Confocal and Airyscan imaging were performed on either a Zeiss LSM 880 or Zeiss LSM 980 microscope. Mounted fillet preparations were imaged with a ×63 oil immersion objective (NA 1.4, DIC; Zeiss) using Zen software (Zeiss Zen Black 2.3 SP1). Airyscan processing of imaging planes and channels was also performed in Zen. Brp puncta surfaces and volumes were calculated using a custom Imaris (Oxford Instruments) routine. We calculated a surface from the SynapGCaMP signal for each NMJ to use as a mask when identifying Brp puncta. Brp puncta were then segmented using the Surfaces function, smoothed with a Gaussian kernel of 0.1 μm, and background subtracted with a 0.15 μm Gaussian kernel within the SynapGCaMP mask. Brp puncta in close proximity to each other were isolated with the Split Touching Objects function with a threshold of 0.15 μm. Low quality ROIs were removed with the Filter Seed Points function (Quality threshold: 59) and applying a minimum volume size (Number of Voxels: 10). To quantify the expression of Unc-13-A and Cac at the AZ, we summed the voxel intensities for these channels within each Brp punctum with the Imaris Spots function. Empirical cumulative distributions of Unc-13-A and Cac intensities were calculated for all Brp puncta for controls and RNAi knockdowns in R (ver 4.1.0).

## Code availability

Code used for all RNA-seq data analysis steps will be available in a Github repository at ^56^ upon publication. QuaSOR custom code is available in the GitHub repository under the filename https://github.com/newmanza/Newman_QuaSOR_2021; https://doi.org/10.5281/zenodo.5711302.

## Data availability

The RNA-seq libraries discussed in this publication and relevant metadata have been deposited in NCBI’s Gene Expression Omnibus^99^ and are accessible through GEO Series GSE250462.

## Acknowledgements

We thank Stephan Sigrist and D.B. Morton for the for the generous gifts of the Unc-13A and Cacophony antibodies. We also thank Nadia Marnani for help with dissections for FACS, Ryan Schultz for guidance on antibody staining, Airy imaging and analysis, and the rest of the Isacoff lab, particularly Ryan Schultz and Adam Hoagland, for helpful discussion. RNAseq was performed in QB3 Genomics, UC Berkeley, Berkeley, CA, RRID:SCR_022170. The study was supported by the National Institutes of Health Grant No. R01NS107506 (E.Y.I.) and a National Science Foundation Graduate Research Fellowship Grant No. DGE 1106400 and DGE 1752814 (C.A.C.).

## Contributions

C.A.C. designed and performed the RNA-seq experiments on *unc13* and *Rbp* RNAi knockdowns, including FACS to isolate type I MNs, RNA-seq library preparation and quality control, and data analysis, as well as Q-PCR validation, with advice from Y.G.C. M.F. performed the FACS to isolate type I MNs and RNA isolation on the *cac* RNAi knockdown and Y.G.C. generated cDNA and quality control, and performed Q-PCR to examine effects on expression level of members of the K_v_ module. D.B. performed the optical quantal imaging of synaptic transmission and QuaSOR analysis with guidance from Z.L.N, who performed preliminary experiments. R.L. performed the antibody staining, Airy scan imaging and analysis. C.A.C. and E.Y.I wrote the paper with input from all of the authors.

