## Supplemental Data for "Deficiency in transmitter release triggers homeostatic transcriptional changes that increase presynaptic excitability"

### Supplementary Material

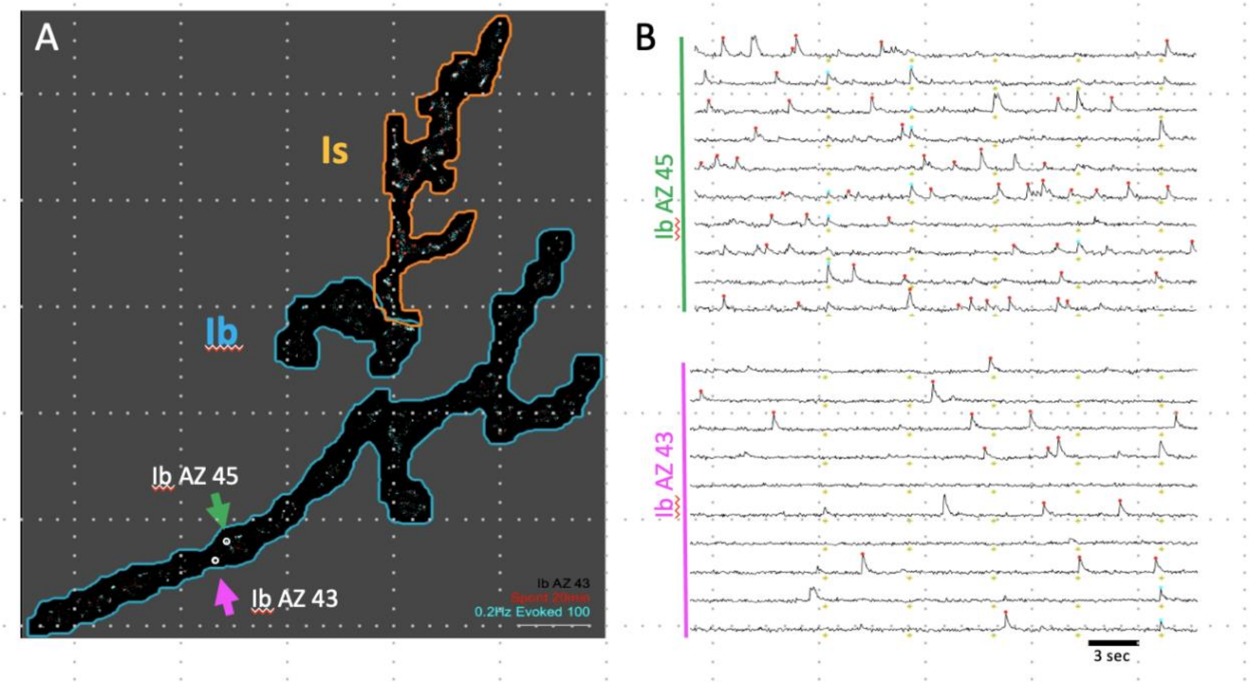


**Figure S1) Quantal imaging of glutamatergic synaptic transmission by Ib motor neurons detected by SynapGCaMP6f in the *Drosophila* larval NMJ.**

**A, B)** Abdominal segment # NMJ showing spontaneous transmission (red) and transmission evoked by action potentials stimulated at 0.2 Hz (blue) by Ib MN# and Is MSNISN to muscle #.

**A)** Image of cumulative transmission.

**B)** Raster plots of transmission events at two identified synapses during first 300 sec period of a movie.

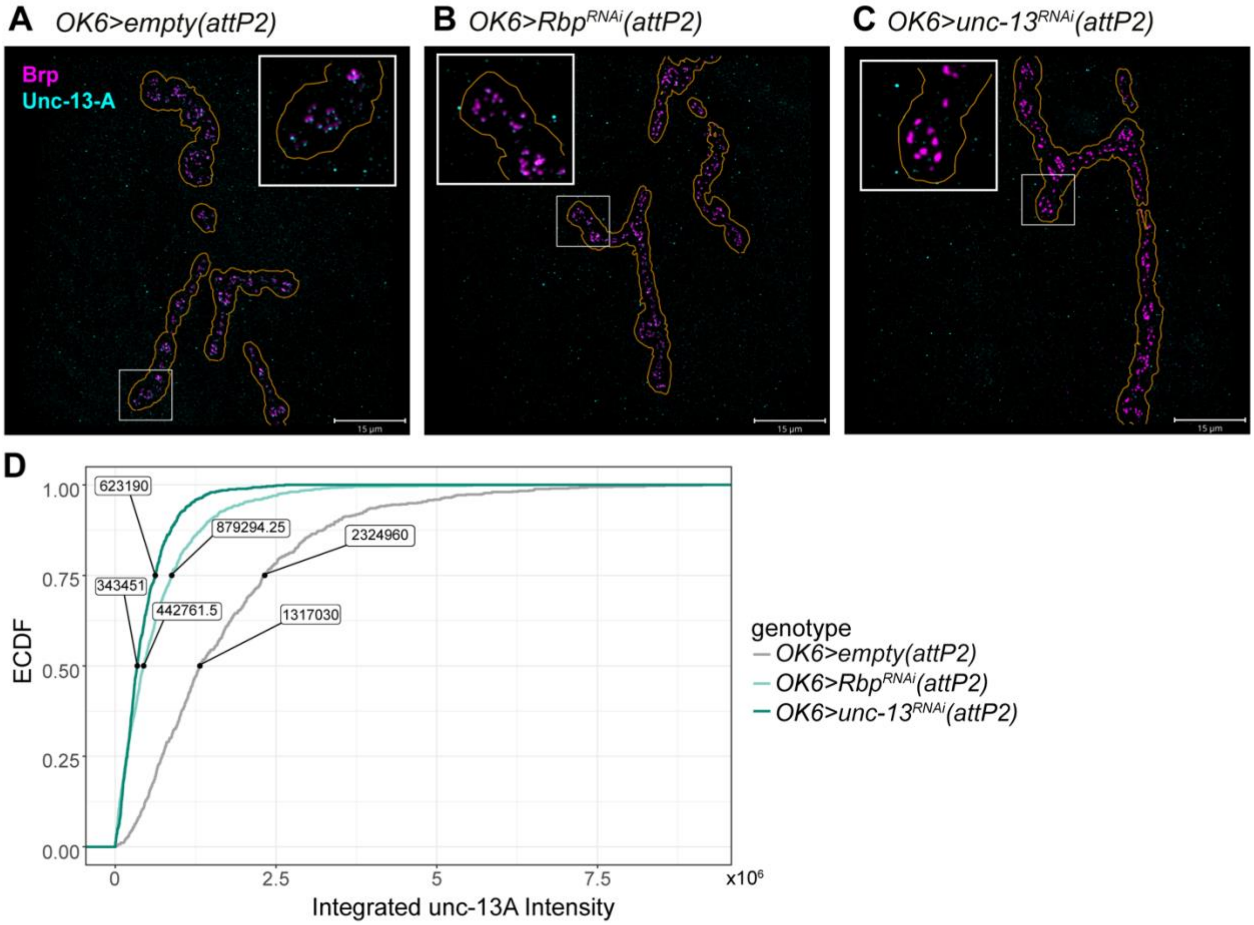


**Figure S2) Rbp and Unc-13 knockdowns decrease Unc-13A at type Ib MN AZs.**

**A-C)** Immunofluorescence stains of Brp (magenta) and Unc-13A (cyan) in type Ib MN terminals (orange outline) of control **(A)** and RNAi knockdowns, *OK6>Rbp$^{RNAi}$* **(B)** and *OK6>unc-13$^{RNAi}$* **(C)**. Scale bar: 15 µm. **D)** Cumulative distribution of summed Unc-13-A voxel intensities in controls and RNAi knockdowns (two-sided KS test, control v *Rbp$^{RNAi}$* *p*-value < 2.2e-16, control vs *unc-13$^{RNAi}$* *p*-value 2.2e-16). Values displayed are the 50$^{th}$ and 75$^{th}$ percentiles for each genotype. (Controls: n = 4 larvae, $n_{Brp\ puncta}$ = 711; *OK6> Rbp$^{RNAi}$*: n = 4 larvae; $n_{Brp\ puncta}$ = 1130; *OK6>unc-13$^{RNAi}$*: n = 4 larvae, $n_{Brp\ puncta}$ = 855).

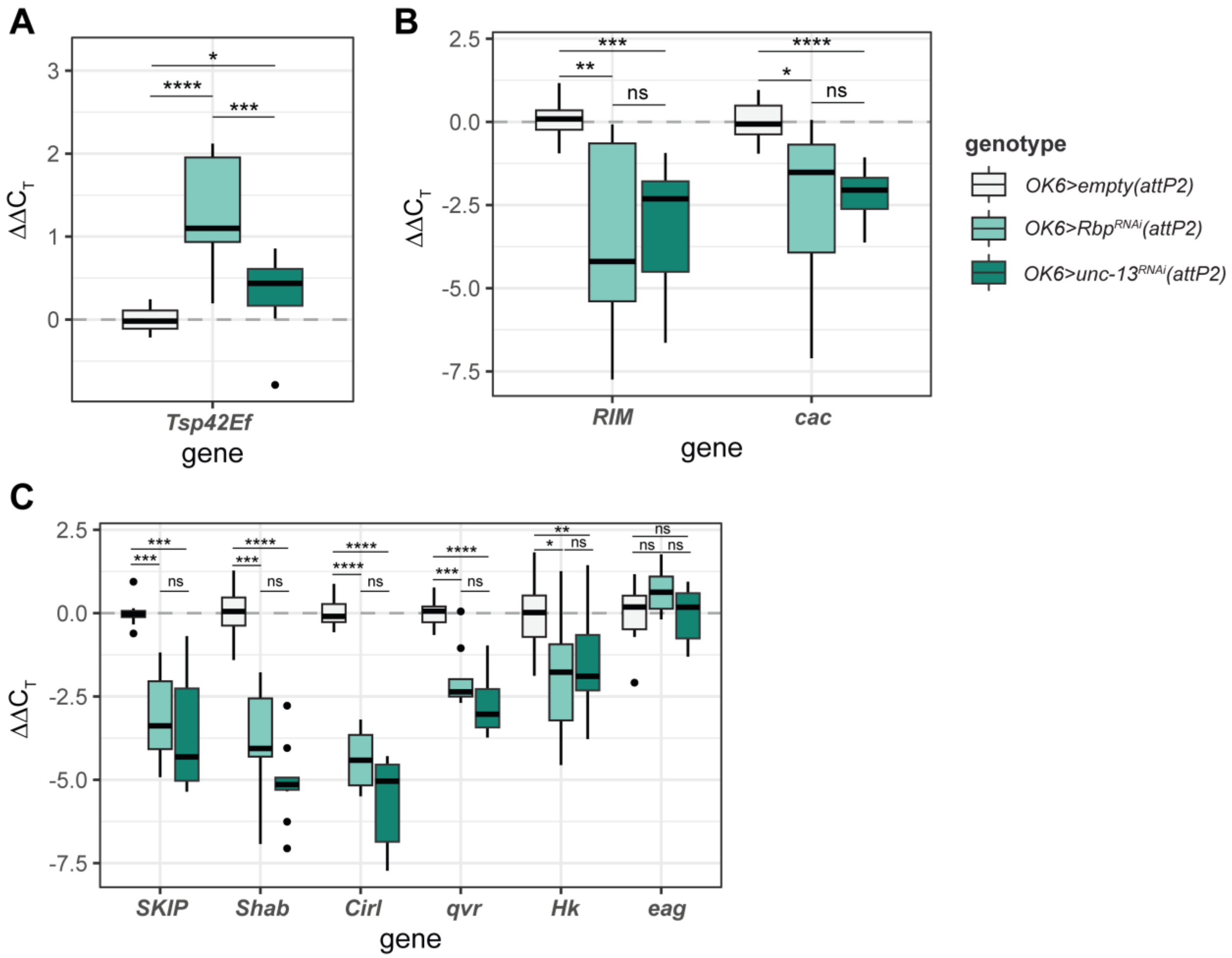


**Figure S3) RT-PCR confirmation of changes in expression of AZ components and Kv channels in RNAi knockdowns.**

**A-C)** $\Delta\Delta C_T$ values of *Tsp42Ef* (**A**), RSSPs (**B**), and Kv channels and their accessory proteins (**C**) in *OK6>Rbp*$^{RNAi}$ and *OK6>unc-13*$^{RNAi}$ (two-tailed T-test *p*-values, ns $p > 0.05$, * $p < 0.01$, ** $p < 0.001$, *** $p < 0.0001$, **** $p <$ 1e-05).

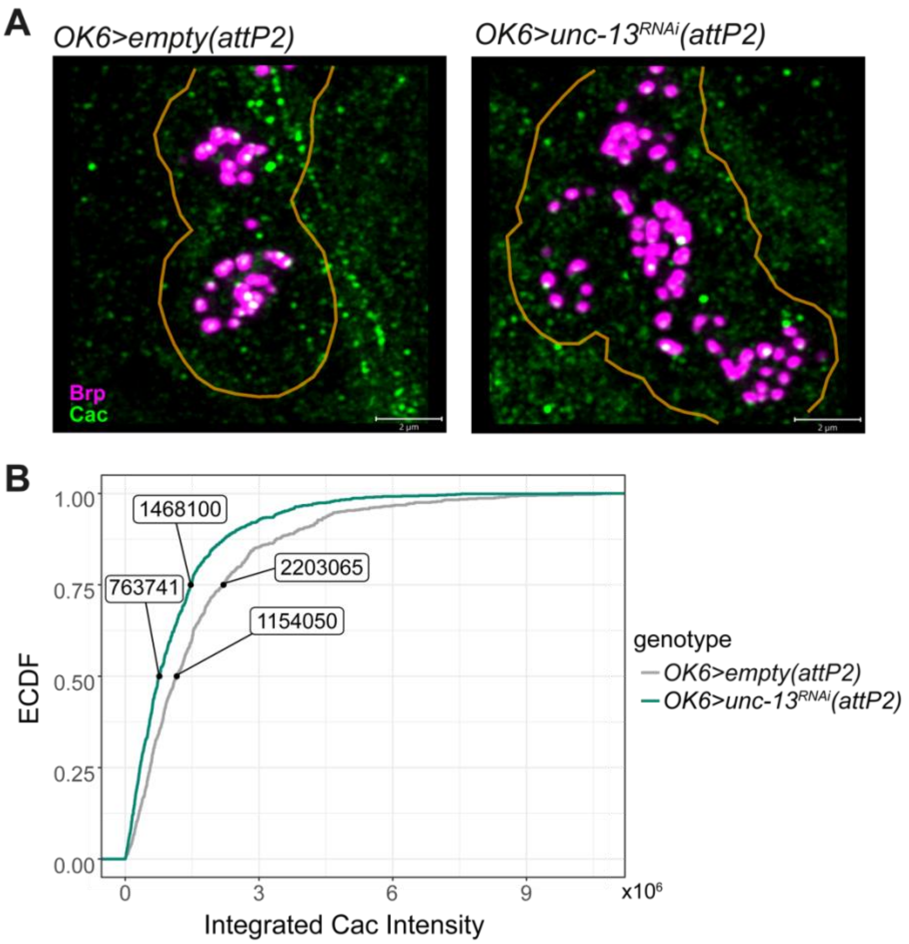


**Figure S4) Unc-13 knockdown decreases Cac at type Ib MN AZs.**

**A)** Immunofluorescence stains of Brp (magenta) and Cac (green) at type Ib MN terminals (orange outline) in controls **(left)** and *OK6>unc-13$^{RNAi}$* **(right)**. Scale bar: 2 µm. **B**) Empirical cumulative distribution of Cac fluorescence within Brp puncta between controls and *OK6>unc-13$^{RNAi}$* (two-sided KS test, p-value = 3.75e-09). Values shown are the 50$^{th}$ and 75$^{th}$ percentiles. (Controls: n = 8 larvae, $n_{Brp\ puncta}$ = 688; *OK6>unc-13$^{RNAi}$*: n = 8 larvae, $n_{Brp\ puncta}$ = 855)

**Supplementary Table 1. Sample information for low-input RNA-seq of control and RNAi type I MNs.** The number of larvae ($n_{VNC}$) and cells ($n_{cells}$) were matched to minimize variability.

| Library | Run | Genotype | $n_{VNCs}$ | $n_{cells}$ |
|---|---|---|---|---|
| OK6_ATTP2_001 | 1 | *OK6>empty(attP2)* | 12 | 6,414 |
| OK6_ATTP2_002 | 1 | *OK6>empty(attP2)* | 12 | 6,414 |
| OK6_ATTP2_003 | 1 | *OK6>empty(attP2)* | 12 | 6,414 |
| OK6_UNC13_001 | 1 | *OK6>unc-13$^{RNAi}$(attP2)* | 19 | 1,753 |
| OK6_UNC13_002 | 1 | *OK6>unc-13$^{RNAi}$(attP2)* | 19 | 1,753 |
| OK6_UNC13_003 | 1 | *OK6>unc-13$^{RNAi}$(attP2)* | 19 | 1,753 |
| OK6_ATTP2_101 | 2 | *OK6>empty(attP2)* | 10 | 3,530 |
| OK6_ATTP2_102 | 2 | *OK6>empty(attP2)* | 10 | 3,530 |
| OK6_ATTP2_103 | 2 | *OK6>empty(attP2)* | 10 | 3,530 |
| OK6_RBP_101 | 2 | *OK6>Rbp$^{RNAi}$(attP2)* | 11 | 3,512 |
| OK6_RBP_102 | 2 | *OK6>Rbp$^{RNAi}$(attP2)* | 11 | 3,512 |
| OK6_RBP_103 | 2 | *OK6>Rbp$^{RNAi}$(attP2)* | 11 | 3,512 |

**Supplementary Table 2. Top 20 up-regulated genes in *OK6>Rbp*$^{RNAi}$ Type I MNs.**

| FlyBase ID | Gene symbol | log-fold-change | *p*-value | Adjusted *p*-value |
|---|---|---|---|---|
| FBgn0038343 | Trissin | 9.33 | 1.29E-12 | 1.77E-10 |
| FBgn0011581 | Ms | 6.71 | 9.39E-50 | 8.92E-46 |
| FBgn0000564 | Eh | 5.64 | 6.69E-05 | 1.05E-03 |
| FBgn0050457 | CG30457 | 5.24 | 1.68E-06 | 4.56E-05 |
| FBgn0053527 | SIFa | 5.14 | 2.96E-05 | 5.29E-04 |
| FBgn0000045 | Act79B | 5.04 | 1.46E-09 | 1.00E-07 |
| FBgn0032096 | Or30a | 5.01 | 3.73E-03 | 2.58E-02 |
| FBgn0000500 | Dsk | 4.44 | 2.00E-03 | 1.58E-02 |
| FBgn0034709 | Swim | 4.35 | 4.67E-03 | 3.06E-02 |
| FBgn0027109 | NPF | 4.05 | 4.81E-36 | 5.08E-33 |
| FBgn0025878 | wrapper | 3.78 | 1.36E-03 | 1.17E-02 |
| FBgn0032895 | twit | 3.06 | 2.57E-32 | 1.88E-29 |
| FBgn0033135 | Tsp42En | 3.06 | 7.06E-06 | 1.60E-04 |
| FBgn0000046 | Act87E | 3.01 | 7.17E-05 | 1.11E-03 |
| FBgn0051370 | CG31370 | 2.99 | 6.00E-03 | 3.71E-02 |
| FBgn0032897 | CG9336 | 2.88 | 3.31E-05 | 5.81E-04 |
| FBgn0023534 | CG17778 | 2.81 | 7.65E-12 | 8.66E-10 |
| FBgn0001258 | Ldh | 2.80 | 2.79E-10 | 2.27E-08 |
| FBgn0034709 | Gsl1 | 2.69 | 1.29E-40 | 4.08E-37 |
| FBgn0034583 | CG10527 | 2.65 | 1.32E-32 | 1.09E-29 |

**Supplementary Table 3. Top 20 up-regulated genes in *OK6>unc-13$^{RNAi}$* Type I MNs.**

| FlyBase ID | Gene symbol | log-fold-change | *p*-value | Adjusted *p*-value |
|---|---|---|---|---|
| FBgn0027109 | NPF | 7.81 | 1.04E-129 | 9.85E-126 |
| FBgn0050457 | CG30457 | 7.20 | 3.66E-11 | 9.38E-09 |
| FBgn0038349 | AOX3 | 7.13 | 1.16E-04 | 4.03E-03 |
| FBgn0032096 | Or30a | 7.08 | 3.72E-05 | 1.58E-03 |
| FBgn0000045 | Act79B | 6.69 | 9.91E-16 | 5.22E-13 |
| FBgn0025878 | wrapper | 6.42 | 3.70E-08 | 4.88E-06 |
| FBgn0036146 | nkt | 6.35 | 4.31E-05 | 1.79E-03 |
| FBgn0039332 | alrm | 6.03 | 1.94E-17 | 1.15E-14 |
| FBgn0028940 | Cyp28a5 | 5.96 | 2.19E-03 | 3.53E-02 |
| FBgn0041087 | wun2 | 5.44 | 2.54E-06 | 1.61E-04 |
| FBgn0038799 | MFS9 | 5.29 | 2.00E-13 | 7.59E-11 |
| FBgn0010389 | htl | 5.20 | 2.13E-03 | 3.47E-02 |
| FBgn0030258 | CG1552 | 5.13 | 1.36E-04 | 4.46E-03 |
| FBgn0033268 | Obp44a | 5.09 | 5.99E-48 | 1.90E-44 |
| FBgn0039915 | Gat | 5.07 | 5.01E-07 | 4.37E-05 |
| FBgn0034588 | CG9394 | 4.99 | 1.99E-03 | 3.29E-02 |
| FBgn0012037 | Ance | 4.94 | 9.81E-04 | 1.93E-02 |
| FBgn0262531 | CG43085 | 4.78 | 1.16E-04 | 4.03E-03 |
| FBgn0030304 | Cyp4g15 | 4.67 | 5.02E-05 | 2.03E-03 |
| FBgn0001145 | Gs2 | 4.63 | 1.45E-17 | 9.17E-15 |

**Supplementary Table 4. Expression of neuropeptide receptor genes in *OK6>Rbp*$^{RNAi}$ Type I MNs.**

| FlyBase ID | Gene symbol | log-fold-change | *p*-value | Adjusted *p*-value |
|---|---|---|---|---|
| FBgn0037408 | NPFR | -1.11 | 1.77E-04 | 3.62E-03 |
| FBgn0004841 | TkR86C | 0.078 | 0.859 | 0.939 |
| FBgn0029723 | Proc-R | -3.71 | 0.088 | 0.253 |
| FBgn0038139 | PK2-R2 | -0.510 | 0.287 | 0.561 |
| FBgn0038140 | PK2-R1 | -1.03 | 6.75E-04 | 9.95E-03 |
| FBgn0038201 | PK1-R | -0.852 | 0.205 | 0.470 |
| FBgn0038874 | ETHR | 0.143 | 0.733 | 0.879 |
| FBgn0038880 | SIFaR | -0.154 | 0.766 | 0.895 |
| FBgn0039396 | CCAP-R | -1.58 | 1.07E-05 | 4.15E-04 |
| FBgn0004842 | RYa-R | -0.573 | 0.291 | 0.566 |
| FBgn0039595 | AstA-R2 | -1.76 | 1.02E-06 | 5.97E-05 |
| FBgn0004622 | TkR99D | 0.184 | 0.490 | 0.732 |
| FBgn0264002 | MsR2 | -0.833 | 0.119 | 0.343 |
| FBgn0035331 | MsR1 | -0.832 | 0.060 | 0.226 |
| FBgn0035385 | FMRFaR | -0.233 | 0.614 | 0.815 |
| FBgn0035610 | Lkr | -0.914 | 0.009 | 0.061 |
| FBgn0053696 | CNMaR | -0.794 | 0.113 | 0.331 |
| FBgn0036278 | CrzR | -0.908 | 0.037 | 0.165 |
| FBgn0036789 | AstC-R2 | -1.86 | 2.35E-07 | 1.85E-05 |
| FBgn0036790 | AstC-R1 | -1.92 | 0.091 | 0.288 |
| FBgn0036934 | sNPF-R | -1.20 | 8.29E-04 | 0.012 |
| FBgn0037100 | CapaR | -0.548 | 0.553 | 0.776 |
| FBgn0033058 | CCHa2-R | -0.067 | 0.784 | 0.904 |
| FBgn0033744 | Dh44-R2 | -0.071 | 0.860 | 0.939 |

| | | | | |
|---|---|---|---|---|
| FBgn0052843 | Dh31-R | -1.38 | 1.63E-04 | 3.39E-03 |
| FBgn0033932 | Dh44-R1 | 0.110 | 0.789 | 0.906 |
| FBgn0050106 | CCHa1-R | -1.29 | 0.017 | 0.096 |
| FBgn0266429 | AstA-R1 | -0.667 | 0.287 | 0.561 |
| FBgn0029768 | SPR | -0.839 | 1.50E-03 | 0.017 |
| FBgn0259231 | CCKLR-17D1 | -0.999 | 0.010 | 0.066 |
| FBgn0030954 | CCKLR-17D3 | -0.708 | 0.168 | 0.421 |
| FBgn0085410 | TrissinR | -1.23 | 2.34E-05 | 7.65E-04 |
| FBgn0003255 | rk | -0.586 | 0.349 | 0.623 |

**Supplementary Table 5. Expression of neuropeptide receptor genes in *OK6>unc-13*$^{RNAi}$ Type I MNs.**

| FlyBase ID | Gene symbol | log-fold-change | *p*-value | Adjusted *p*-value |
|---|---|---|---|---|
| FBgn0037408 | NPFR | -1.0766581 | 3.49E-04 | 0.014 |
| FBgn0004841 | TkR86C | 0.57818307 | 0.192 | 0.587 |
| FBgn0029723 | Proc-R | 1.46 | 0.443 | 0.745 |
| FBgn0038139 | PK2-R2 | -0.5383246 | 0.270 | 0.667 |
| FBgn0038140 | PK2-R1 | -0.8116457 | 9.61E-03 | 0.128 |
| FBgn0038201 | PK1-R | -0.4089712 | 0.551 | 0.860 |
| FBgn0038874 | ETHR | 0.71183909 | 0.093 | 0.436 |
| FBgn0038880 | SIFaR | -3.2700512 | 7.68E-08 | 1.42E-05 |
| FBgn0039396 | CCAP-R | -1.1086678 | 2.97E-03 | 0.062 |
| FBgn0004842 | RYa-R | 0.03736253 | 0.946 | 0.987 |
| FBgn0039595 | AstA-R2 | -0.8153448 | 0.025 | 0.221 |
| FBgn0004622 | TkR99D | 0.75852444 | 4.84E-03 | 0.084 |
| FBgn0264002 | MsR2 | -0.2289584 | 0.674 | 0.911 |
| FBgn0035331 | MsR1 | -0.2848718 | 0.523 | 0.845 |
| FBgn0035385 | FMRFaR | 0.05374804 | 0.909 | 0.977 |
| FBgn0035610 | Lkr | -0.4722641 | 0.183 | 0.576 |
| FBgn0053696 | CNMaR | 0.31123026 | 0.536 | 0.853 |
| FBgn0036278 | CrzR | 0.10520534 | 0.810 | 0.949 |
| FBgn0036789 | AstC-R2 | -1.418269 | 1.00E-04 | 5.36E-03 |
| FBgn0036790 | AstC-R1 | -0.839652 | 0.466 | 0.814 |
| FBgn0036934 | sNPF-R | -0.4859298 | 0.178 | 0.571 |
| FBgn0037100 | CapaR | 0.91923605 | 0.320 | 0.707 |
| FBgn0033058 | CCHa2-R | 0.03495696 | 0.888 | 0.973 |
| FBgn0033744 | Dh44-R2 | -0.6429156 | 0.144 | 0.524 |

| | | | | |
|---|---|---|---|---|
| FBgn0052843 | Dh31-R | 0.09442688 | 0.797 | 0.945 |
| FBgn0033932 | Dh44-R1 | -0.3911786 | 0.348 | 0.729 |
| FBgn0050106 | CCHa1-R | -0.8504323 | 0.124 | 0.493 |
| FBgn0266429 | AstA-R1 | -1.1907176 | 0.059 | 0.351 |
| FBgn0029768 | SPR | -1.5134016 | 5.36E-08 | 1.04E-05 |
| FBgn0259231 | CCKLR-17D1 | 0.13379524 | 0.732 | 0.930 |
| FBgn0030954 | CCKLR-17D3 | -0.0631631 | 0.902 | 0.977 |
| FBgn0085410 | TrissinR | -0.6916678 | 0.021 | 0.201 |
| FBgn0003255 | rk | -0.9592637 | 0.150 | 0.534 |

**Supplementary Table 6. Expression of ionotropic receptor genes in *OK6>Rbp*$^{RNAi}$ Type I MNs.**

| **Cholinergic Receptors** | | | | |
|---|---|---|---|---|
| FlyBase ID | Gene symbol | log-fold-change | *p*-value | Adjusted *p*-value |
| FBgn0032151 | nAChRalpha6 | -1.27 | 4.54E-08 | 2.00E-06 |
| FBgn0000036 | nAChRalpha1 | -1.25 | 4.94E-04 | 5.46E-03 |
| FBgn0028875 | nAChRalpha5 | -0.997 | 4.52E-04 | 5.06E-03 |
| FBgn0015519 | nAChRalpha3 | -0.369 | 0.402 | 0.630 |
| FBgn0266347 | nAChRalpha4 | -0.298 | 0.215 | 0.447 |
| FBgn0086778 | nAChRalpha7 | 0.014 | 0.945 | 0.977 |
| FBgn0004118 | nAChRbeta2 | 0.484 | 8.17E-02 | 0.242 |
| FBgn0000039 | nAChRalpha2 | 0.530 | 7.37E-03 | 4.31E-02 |
| FBgn0000038 | nAChRbeta1 | 0.546 | 3.92E-02 | 0.147 |
| **GABAergic Receptors** | | | | |
| FlyBase ID | Gene symbol | log-fold-change | *p*-value | Adjusted *p*-value |
| FBgn0001134 | Grd | -3.02 | 0.131 | 0.330 |
| FBgn0033558 | CG12344 | -1.80 | 9.38E-02 | 0.265 |
| FBgn0030707 | CG8916 | -0.814 | 0.605 | 0.788 |
| FBgn0004244 | Rdl | -0.721 | 9.13E-04 | 8.70E-03 |
| FBgn0010240 | Lcch3 | 0.135 | 0.632 | 0.806 |
| **Glutamatergic Receptors** | | | | |
| FlyBase ID | Gene symbol | log-fold-change | *p*-value | Adjusted *p*-value |
| FBgn0039916 | Ekar | -3.81 | 4.17E-04 | 4.74E-03 |
| FBgn0051201 | GluRIIE | -1.75 | 5.38E-02 | 0.183 |
| FBgn0264000 | GluRIB | -0.94 | 1.63E-05 | 3.23E-04 |
| FBgn0024963 | GluCalpha | -0.63 | 2.69E-03 | 1.99E-02 |

| | | | | |
|---|---|---|---|---|
| FBgn0004619 | GluRIA | -0.33 | 0.201 | 0.428 |
| FBgn0039927 | CG11155 | -0.27 | 0.168 | 0.383 |
| FBgn0053513 | Nmdar2 | -0.188 | 0.371 | 0.602 |
| FBgn0010399 | Nmdar1 | 0.066 | 0.721 | 0.860 |
| FBgn0038837 | KaiR1D | 0.26 | 0.290 | 0.528 |
| FBgn0038840 | Grik | 0.55 | 0.634 | 0.808 |

**Supplementary Table 7. Expression of ionotropic receptor genes in *OK6>unc-13$^{RNAi}$* Type I MNs.**

| **Cholinergic Receptors** | | | | |
|---|---|---|---|---|
| FlyBase ID | Gene symbol | log-fold-change | *p*-value | Adjusted *p*-value |
| FBgn0032151 | nAChRalpha6 | -1.34 | 1.15E-08 | 1.80E-06 |
| FBgn0015519 | nAChRalpha3 | -1.02 | 2.22E-02 | 0.162 |
| FBgn0000036 | nAChRalpha1 | -0.789 | 2.84E-02 | 0.188 |
| FBgn0028875 | nAChRalpha5 | -0.648 | 0.234 | 0.167 |
| FBgn0086778 | nAChRalpha7 | -0.428 | 4.21E-02 | 0.237 |
| FBgn0266347 | nAChRalpha4 | -0.199 | 0.416 | 0.725 |
| FBgn0000039 | nAChRalpha2 | -0.170 | 3.96E-01 | 0.708 |
| FBgn0004118 | nAChRbeta2 | 0.046 | 0.870 | 0.953 |
| FBgn0000038 | nAChRbeta1 | 0.118 | 0.657 | 0.868 |
| **GABAergic Receptors** | | | | |
| FlyBase ID | Gene symbol | log-fold-change | *p*-value | Adjusted *p*-value |
| FBgn0001134 | Grd | -4.25 | 5.31E-02 | 0.271 |
| FBgn0004244 | Rdl | -1.13 | 1.93E-07 | 1.95E-05 |
| FBgn0010240 | Lcch3 | 0.310 | 0.278 | 0.607 |
| FBgn0030707 | CG8916 | 0.214 | 0.894 | 0.964 |
| FBgn0033558 | CG12344 | 0.223 | 0.833 | 0.939 |
| **Glutamatergic Receptors** | | | | |
| FlyBase ID | Gene symbol | log-fold-change | *p*-value | Adjusted *p*-value |
| FBgn0039916 | Ekar | -1.78 | 8.60E-02 | 0.348 |
| FBgn0024963 | GluCalpha | -1.01 | 1.91E-06 | 1.26E-04 |
| FBgn0264000 | GluRIB | -0.87 | 1.01E-04 | 3.60E-03 |
| FBgn0039927 | CG11155 | -0.52 | 8.76E-03 | 8.99E-02 |

| | | | | |
|---|---|---|---|---|
| FBgn0004619 | GluRIA | -0.43 | 0.105 | 0.384 |
| FBgn0053513 | Nmdar2 | -0.29 | 0.168 | 0.483 |
| FBgn0051201 | GluRIIE | -0.21 | 0.814 | 0.930 |
| FBgn0038837 | KaiR1D | 0.01 | 0.974 | 0.991 |
| FBgn0010399 | Nmdar1 | 0.49 | 0.957 | 9.57E-02 |
| FBgn0038840 | Grik | 0.61 | 0.608 | 0.843 |
